# Electrostatically regulated active site assembly governs reactivity in non-heme iron halogenases

**DOI:** 10.1101/2023.05.25.542349

**Authors:** Elizabeth R. Smithwick, R. Hunter Wilson, Sourav Chatterjee, Yu Pu, Joseph J. Dalluge, Anoop Rama Damodaran, Ambika Bhagi-Damodaran

## Abstract

Non-heme iron halogenases (NHFe-Hals) catalyze the direct insertion of a chloride/bromide ion at an unactivated carbon position using a high-valent haloferryl intermediate. Despite more than a decade of structural and mechanistic characterization, how NHFe-Hals preferentially bind specific anions and substrates for C-H functionalization remains unknown. Herein, using lysine halogenating BesD and HalB enzymes as model systems, we demonstrate strong positive cooperativity between anion and substrate binding to the catalytic pocket. Detailed computational investigations indicate that a negatively charged glutamate hydrogen-bonded to iron’s equatorial-aqua ligand acts as an electrostatic lock preventing both lysine and anion binding in the absence of the other. Using a combination of UV-Vis spectroscopy, binding affinity studies, stopped-flow kinetics investigations, and biochemical assays, we explore the implication of such active site assembly towards chlorination, bromination, and azidation reactivities. Overall, our work highlights previously unknown features regarding how anion-substrate pair binding govern reactivity of iron halogenases that are crucial for engineering next-generation C-H functionalization biocatalysts.

---

The use of enzymes as industrial catalysts has gained increasing attention in recent years for their ability to achieve complex chemical transformations with a high degree of regio- and stereo-selectivity under environment-friendly conditions.^1–6^ Non-heme iron halogenases (NHFe-Hals), in particular, hold great value as biocatalytic platforms as they are able to install a variety of functional moieties (Cl^-^, Br^-^, N_3_ ^-^, NO_2_^-^) at an unactivated *sp*^3^-hybridized carbon in a selective manner.^7–12^ However, only a handful of identified NHFe-Hals are able to act on freestanding substrates, limiting their substrate scope currently to some amino acids,^7^ alkaloids,^12,13^ and nucleotides^14,15^ (**Fig. 1A**). A frequently used strategy to counteract the limited substrate scope of NHFe-Hals has been to engineer a related class of better characterized enzymes called non-heme iron hydroxylases (NHFe-Hyds) which offer a broader range of substrates.^16–18^ NHFe-Hals and NHFe-Hyds differ in their primary coordination sphere (PCS) seemingly by a single amino acid substitution, with NHFe-Hyds featuring a canonical 2-His-1-carboxylate facial triad that coordinates the iron center.^19^ In NHFe-Hals, however, the carboxylate of the facial triad is substituted for a glycine or alanine, allowing an open coordination position for chloride to bind.^20^ Initial attempts to reprogram NHFe-Hyds into halogenases by this single amino acid substitution of Glu/Asp to Ala/Gly were unsuccessful, indicating a more complex molecular mechanism at play.^21,22^ To expand the reactivity of homologous hydroxylases, as well as to promote non-native C-H functionalization catalysis, a rigorous understanding of how NHFe-Hals facilitate binding, activation, and reactivity with their substrates and anions is needed.

**Figure 1.**
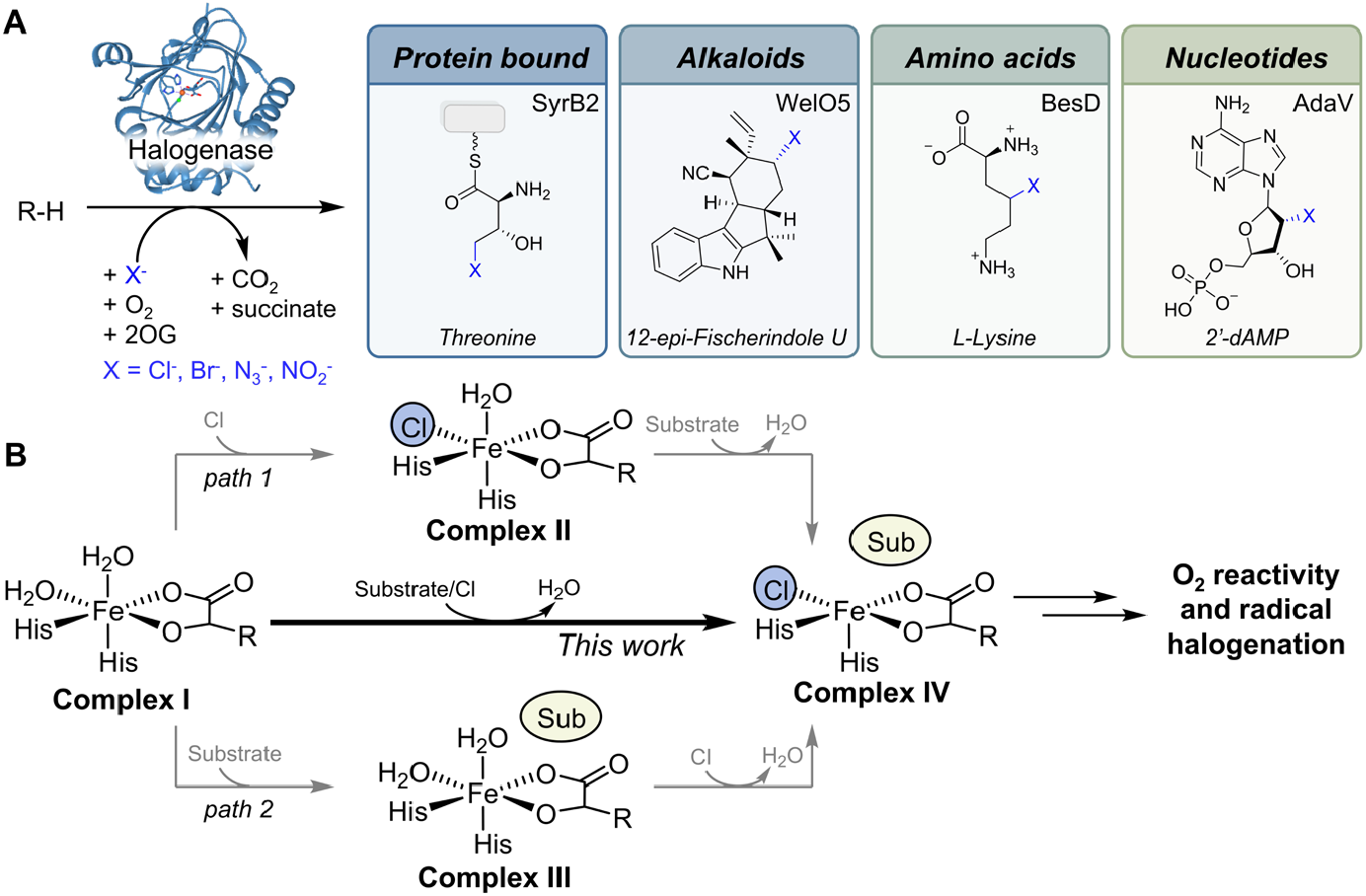
**(A)** Simplified halogenation reaction and current substrate scope of NHFe-Hals with specific examples of each substrate category. **(B)** Schematic showing complexes formed to significant levels in NHFe-Hals at physiological concentrations of chloride and substrate.

In the absence of substrate and anion, the iron center of NHFe-Hals is six-coordinate (*6c*) with two histidines, two aqua ligands, and 2-oxoglutarate (2OG) composing the PCS (complex **I** in **Fig. 1B**). Previous studies have found that the unreactive complex **I** is converted to the catalytically relevant complex **IV** in a random sequential order of anion and substrate binding events (path 1 or 2, **Fig. 1B)**. ^23–29^ Once formed, the substrate/chloride bound complex **IV** will react rapidly with O_2_ to form a high-valent haloferryl intermediate that abstracts a hydrogen atom from the substrate.^30^ The newly formed substrate radical can then rebound with iron-coordinated chloride to form the chlorinated product.^31,32^ Here, we demonstrate that lysine-halogenating enzymes BesD and HalB exhibit a strong positive heterotropic cooperativity between lysine and chloride binding. Through molecular dynamics (MD) simulations and density functional theory (DFT) calculations, we find that a negatively charged residue E119 in BesD locks the active site inhibiting both lysine and anion binding in the absence of the other. Only when chloride/bromide/azide anions work in tandem with lysine can they both effectively bind in the catalytic pocket enabling respective reactivity. Overall, we demonstrate how electrostatically regulated active site assembly in NHFe-Hals tightly controls its C-H functionalization reactivity and prevents unproductive reaction cycling to reduce off-target reactions.

To understand how lysine and chloride bind to BesD, we monitored halogenase active site assembly using UV-Vis spectroscopy. NHFe-2OG dependent enzymes exhibit a broad metal-to-ligand charge transfer (MLCT) band between the Fe^II^ center and the bidentate coordinated 2OG ligand centered around 475-550 nm.^33^ This spectral feature is sensitive to changes in the PCS of Fe and has been used to study substrate binding in NHFe-Hyds and anion binding in NHFe-Hals.^11,27,33–37^ Upon the addition of Fe^II^ and 2OG to BesD, an MLCT band forms with a peak maximum centered at 480 nm (**Fig. 2A** grey spectrum) that can be assigned to a *6c*-diaqua Fe species (**I**), similar to the complex previously characterized in SyrB2.^11^ However, unlike SyrB2,^11,37^ the peak maximum of the MLCT band remains mostly unperturbed upon the addition of chloride to the BesD-Fe-2OG complex suggesting that the chloride is not able coordinate to Fe^II^, even at >4.5 mM excess concentrations (**Fig. 2A** purple spectrum). It is not until lysine is added to the Cl^-^ containing mixture that a distinctive shift in peak maximum to 500 nm is observed (**Fig. 2A** blue spectrum), indicating that a change in the PCS of iron has occurred. After reversing the order of chloride and lysine additions, we now find that the addition of only lysine (>4.5 mM excess) induce hardly any spectral shift, in contrast to other characterized NHFe enzymes.^27,28,33^ Only after the addition of chloride to the BesD-Fe-2OG-lysine mixture does the peak maximum shift indicating that both the substrate and chloride work in tandem to alter the PCS of Fe^II^ (**Fig. 2B**). To investigate the catalytic relevance of complex **I** and various lysine- and chloride-added forms of BesD, we employed stopped-flow UV-Vis spectroscopy^38–40^ to explore their O_2_ reactivity profiles by monitoring the formation of the high-valent ferryl intermediate (Fe^IV^=O) through transient absorbance increases at ∼320 nm as has been established previously.^11,25,34^ Analogous to these studies, we observe a rapid increase in the absorbance at 320 nm when the BesD-Fe-2OG-Lys-Cl complex is mixed with O_2_-saturated buffer (∼1 mM), indicating an efficient reaction with oxygen (**Fig. 2C** blue curve). Based on such a reactivity profile, we assign the BesD-Fe-2OG-Lys-Cl complex with MLCT centered at 500 nm (**Fig. 2A-B**, blue spectrum) to be the catalytically primed *5c*-chloro Fe complex **IV**. In contrast, if either chloride and/or lysine are absent, the protein complex exhibits negligible reactivity with O_2_ (**Fig. 2C** grey/purple/green spectrum). The minor initial reactivity observed in the lysine-only condition is likely due to chloride contamination which has been mentioned as an issue in previous studies.^8,11^ In all, UV-Vis studies combined with stopped-flow kinetic investigations reveal that both lysine and chloride cooperatively bind to convert complex **I** to catalytically primed complex **IV** in BesD (**Fig. 2D**). We note that the peak shift in the MLCT band upon simultaneous addition of lysine and chloride represents such a conversion (**Fig. 2A-B**). Furthermore, we observe similar trends in HalB (**Fig. S1**) suggesting that this effect could be a common mechanism across this family of NHFe-Hals.

**Figure 2.**
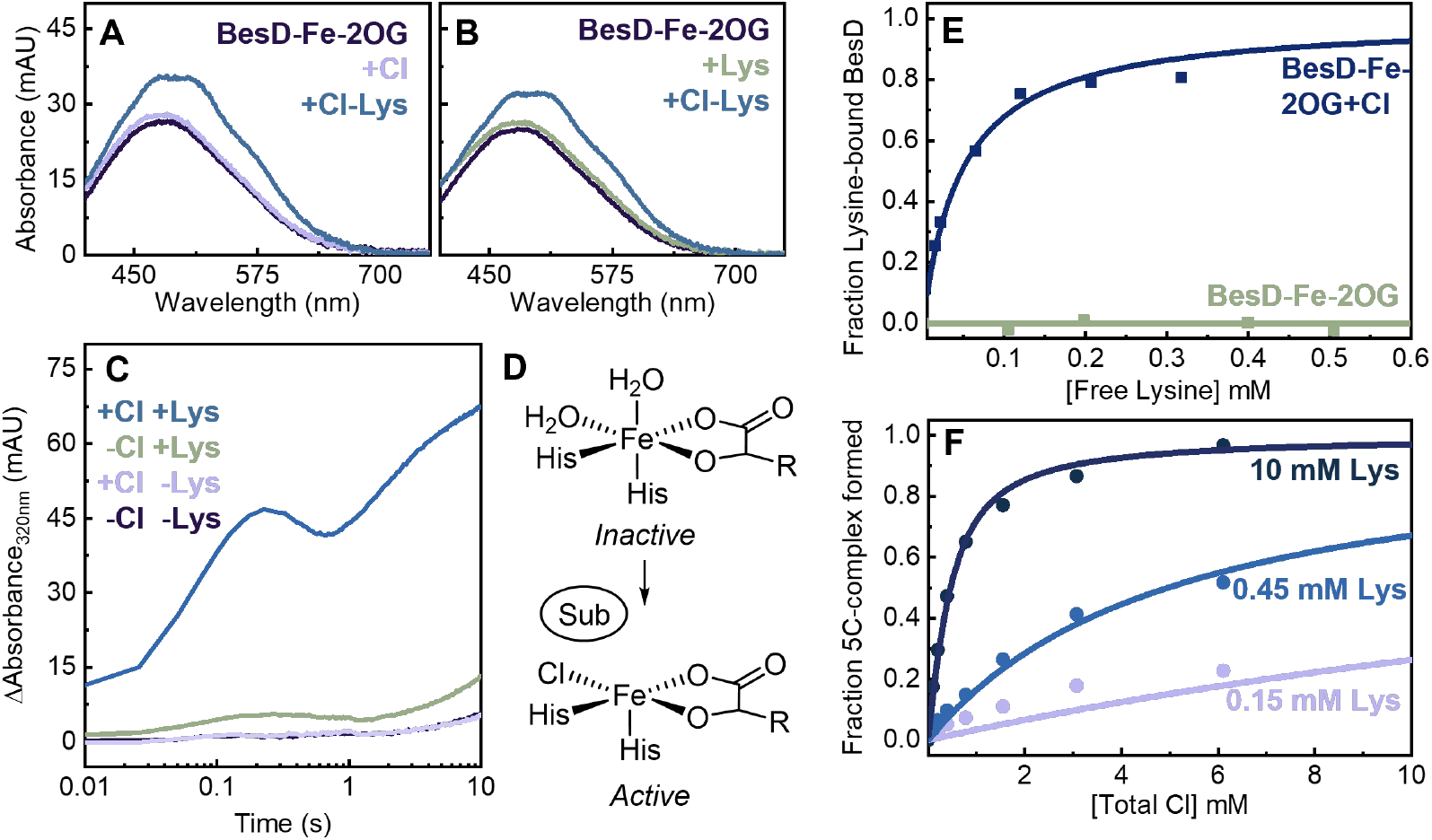
Spectral changes in MLCT band of BesD-Fe-2OG complex upon sequential additions of **(A)** Cl then lysine or **(B)** lysine then Cl. (**C**) Increase in absorbance at 320 nm over time as observed by stopped-flow UV-Vis spectroscopic kinetics for various combinations of lysine/chloride. (**D**) Conversion of **I** to catalytically primed **IV** in BesD upon cooperative binding of lysine and chloride. (**E**) Equilibrium dialysis-based binding studies of lysine to BesD-Fe-2OG in the presence and absence of chloride. (**F**) Titration curves for chloride in BesD at different lysine concentrations where the increasing absorbance of the 518 nm species has been converted to fraction of *5c*-species formed.

After uncovering this substrate-anion relationship, we conducted binding affinity studies to explore the nature of their association and its impact on the formation of the primed complex **IV** in BesD. Equilibrium dialysis of varied concentrations of lysine against BesD and subsequent HPLC analysis of free-lysine in the protein-free compartment revealed a strong chloride-dependent lysine binding behavior (**Fig. S2-S3**). In the absence of chloride, we observe that lysine does not show any measurable association to BesD even at 0.5 mM concentrations (**Fig. 2E**, green curve). On the other hand, the apparent binding affinity of lysine is strongly enhanced (*K*_d,app_ = 50 µM) in the presence of 30 mM chloride (**Fig. 2E**, blue curve). Next, we titrated chloride to the BesD-Fe-2OG complex in the presence of various lysine concentrations and employed UV-Vis spectroscopy to probe the formation of the *5c*-species as a measure of chloride binding (**Fig. S4**). Here, we again observe chloride binding to BesD has a strong lysine-concentration dependence. We measure apparent chloride binding affinities of 28, 4.8, and 0.29 mM in the presence of 0.15, 0.45, and 10 mM lysine, respectively (**Fig. 2F**). Taken together, these chloride/lysine binding affinity studies clearly demonstrate a strong positive cooperativity that governs the assembly of the BesD active site.

To probe the molecular basis of this cooperativity, we performed MD simulations on different BesD active site configurations. In simulations with the *6c*-diaqua BesD-Fe complex **I**, we observe a strong hydrogen bonding (H-bonding) network composed of E119 and W137 that stabilizes the equatorial-aqua ligand (**Fig. 3A**). Both residues are highly conserved in the BesD family of halogenases and previous work shows that mutating E119 to Ala results in a loss of activity.^7^ Simulations of complex **IV** in BesD reveals a dramatic reorganization of this H-bond network (**Fig. 3B**). Comparing the two configurations, we observe that lysine forms a salt-bridge with the E119 residue which necessitates disruption of the rather strong H-bond (sampled over 94±3% of the simulation) between E119 and iron’s equatorial-aqua ligand creating an energy barrier to lysine binding. The presence of chloride at the equatorial site will not only help disrupt these H-bonds, but also provide favorable electrostatic interactions for lysine binding. Similarly, the negative charge on E119 near the active site would repel chloride until screened by positively charged lysine. By generating electrostatic potential maps based on optimized geometries of MD simulations, we can see that the presence of E119 creates a negative potential at the equatorial site in the absence of lysine as indicated by the red color of the electrostatic surface (**Fig. 3C**). This would make anion binding electrostatically unfavorable. Alternatively, when the positively charged lysine sidechain is present, the electrostatic environment becomes far more positive and therefore more conducive to anion binding (**Figs. 3D, S5**). Overall, the negatively charged E119 residue H-bonded to iron’s equatorial-aqua ligand of complex **I** acts as an electrostatic lock that inhibits chloride and lysine binding in the absence of the other. When lysine and chloride are both present, they disrupt the E119-aqua interaction and push E119 away from the Fe^II^ center (**Fig. 3E**) transforming the electrostatic environment in the active site. These studies provide insights into the experimentally observed positive cooperativity in lysine-chloride pair binding and reveal the role of E119 as an electrostatic lock in BesD.

**Figure 3.**
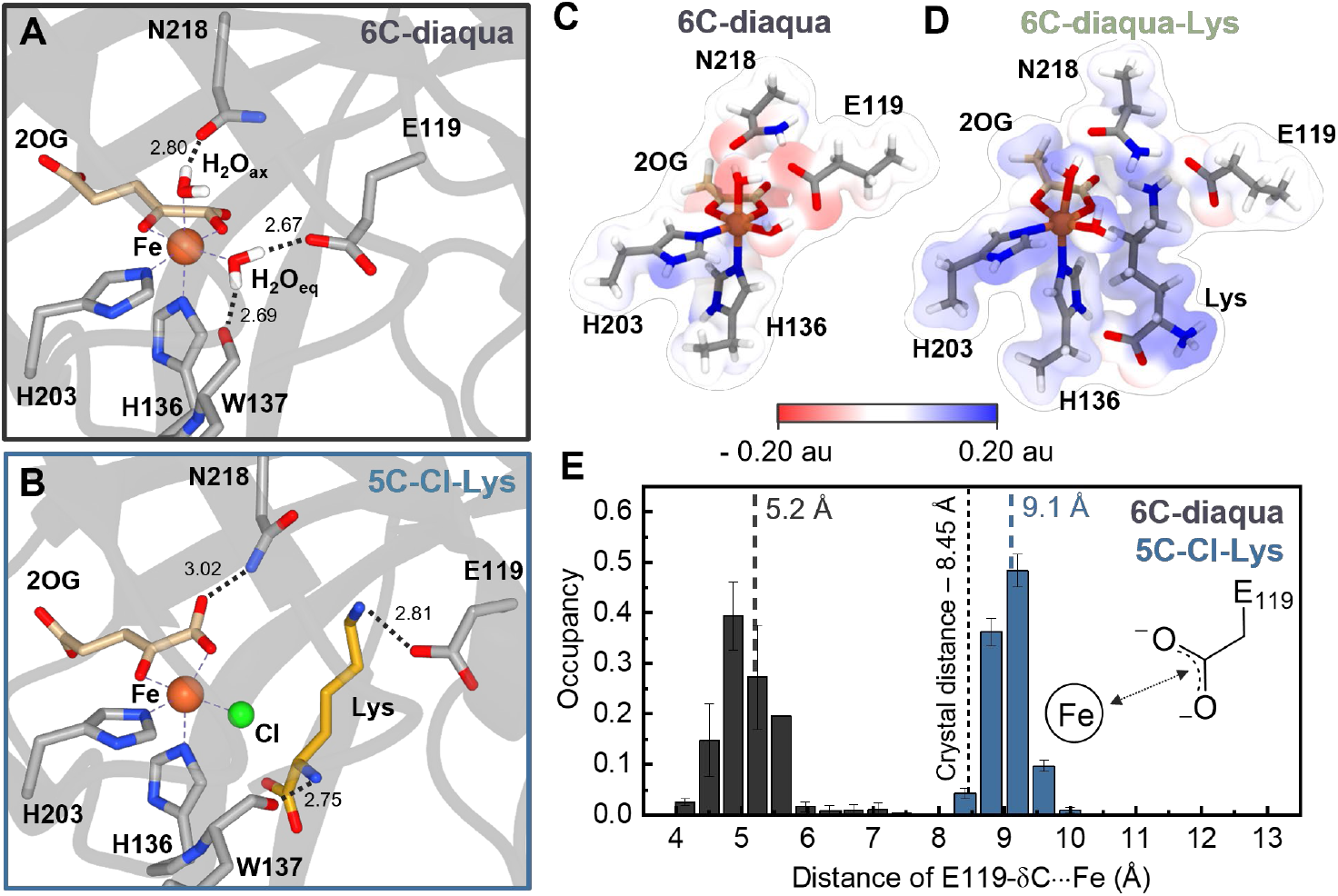
Representative structures of MD simulated (**A**) complex **I** and (**B**) complex **IV** in BesD. Electrostatic surfaces calculated based on calculated RESP charges of optimized representative snapshots of the (**C**) complex **I** in BesD and (**D**) lysine-bound BesD-Fe-2OG. (**E**) Histogram of distance between iron and the δ-carbon of E119 achieved during the simulation duplicates of the different active site configurations. Dotted lines represent the median distance averaged between duplicate simulations.

We also explored BesD’s reactivity profile with other anions to better understand the implications of such complex active site assembly on C-H functionalization.^7–12^ We scanned the ability of BesD to utilize non-native anions with UPLC-MRM-MS/MS-based reaction studies which revealed that while BesD could azidate and brominate lysine (confirmed by accurate mass measurements, **Table S2**), no evidence of nitration was detected (**Figs. 4A-C, S6-S8**). Like chloride, perturbations of the MLCT band with azide and bromide are minimal until the addition of lysine, indicating lysine-dependent binding is operant with non-native anions as well (**Fig. 4D-F**). Aligning with the reactivity assays, nitrite failed to produce any definitive peak change beyond its own intrinsic absorbance pattern. Overall, these studies confirm that the reactivity profile of BesD is controlled by the active-site assembly of substrate-anion pairs and could be an important factor in future enzyme engineering endeavors.

**Figure 4.**
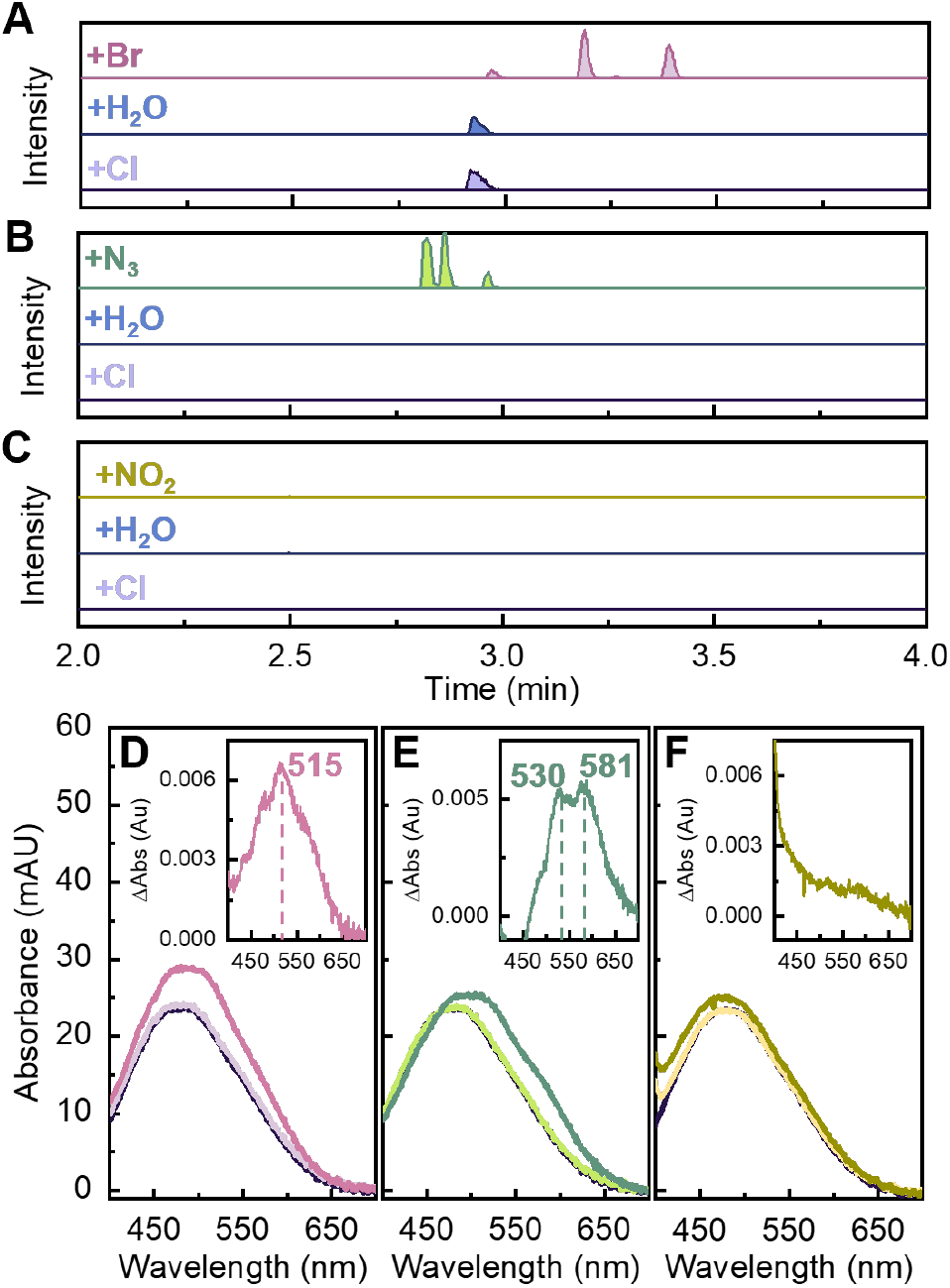
UPLC-MS/MS chromatograms of BesD reactions with non-native anions: (**A**) Br^-^, (**B**) N_3_ ^-^, and (**C**) NO_2_^-^. Anaerobic UV-Vis spectra of anion additions with (**D**) Br^-^, (**E**) N_3_ ^-^, and (**F**) NO_2_^-^ and subsequent lysine additions to BesD-Fe-2OG complexes. Inset shows difference spectrum for each panel.

In summary, this work establishes electrostatically regulated cooperative binding of anion-substrate pair as a new form of reactivity control in BesD and HalB. Such controls in BesD and other similar halogenases likely leads to greater conservation of cellular resources as other halogenases can experience an uncoupled consumption of O_2_ in the chloro-bound form.^24^ Understanding if other freestanding NHFe-Hals adhere to similar or divergent mechanisms of control would enable substrate scope expansion in NHFe-Hals as well as facilitate the conversion of NHFe-Hyds to halogenases.

## ASSOCIATED CONTENT

### Supporting Information

Experimental details on Protein Purification/Expression, Equilibrium UV-Vis studies, Equilibrium Dialysis Studies, Stopped-flow UV/Vis Spectroscopy, Molecular Dynamics, DFT calculations, Activity Assays, and Mass Spectrometry can be found in supporting information. This material is available free of charge via the Internet at http://pubs.acs.org

## AUTHOR INFORMATION

### Notes

The authors declare no competing financial interests.

## ACKNOWLEDGMENT

ERS and RHW acknowledge the support of the National Institute of Health Chemical Biology Training Grant (T32GM132029). The halogenation part of this work was supported by NSF CBET (grant # 2046527). The azidation/nitration part of this work was supported by the Regents of the University of Minnesota. The authors would like to thank Prof. Michelle Chang (U.C. Berkeley) for BesD and HalB plasmids. All color palettes for figures were taken from Ref. 41.

## REFERENCES

(1) Bornscheuer, U. T.; Huisman, G. W.; Kazlauskas, R. J.; Lutz, S.; Moore, J. C.; Robins, K. Engineering the Third Wave of Biocatalysis. Nature 2012. https://doi.org/10.1038/nature11117.

(2) Hanefeld, U.; Hollmann, F.; Paul, C. E. Biocatalysis Making Waves in Organic Chemistry. Chem. Soc. Rev. 2022. https://doi.org/10.1039/d1cs00100k.

(3) Chapman, J.; Ismail, A. E.; Dinu, C. Z. Industrial Applications of Enzymes: Recent Advances, Techniques, and Outlooks. Catalysts 2018, 8 (6), 20–29. https://doi.org/10.3390/catal8060238.

(4) Tian, J.; Garcia, A. A.; Donnan, P. H.; Bridwell-rabb, J. Leveraging a Structural Blueprint to Rationally Engineer the Rieske Oxygenase TsaM. Biochemistry 2023. https://doi.org/10.1021/acs.biochem.3c00150.

(5) Wang, Y.; Davis, I.; Shin, I.; Xu, H.; Liu, A. Molecular Rationale for Partitioning between C-H and C-F Bond Activation in Heme-Dependent Tyrosine Hydroxylase. J. Am. Chem. Soc. 2021, 143 (12), 4680–4693. https://doi.org/10.1021/jacs.1c00175.

(6) Lewis, J. C.; Coelho, P. S.; Arnold, F. H. Enzymatic Functionalization of Carbon–Hydrogen Bonds. Chem. Soc. Rev. 2011, 40 (4), 2003–2021. https://doi.org/10.1039/c0cs00067a.

(7) Neugebauer, M. E.; Sumida, K. H.; Pelton, J. G.; McMurry, J. L.; Marchand, J. A.; Chang, M. C. Y. A Family of Radical Halogenases for the Engineering of Amino-Acid-Based Products. Nat. Chem. Biol. 2019, 15 (10), 1009–1016. https://doi.org/10.1038/s41589-019-0355-x.

(8) Zhu, Q.; Hillwig, M. L.; Doi, Y.; Liu, X. Aliphatic Halogenase Enables Late-Stage C-H Functionalization: Selective Synthesis of a Brominated Fischerindole Alkaloid with Enhanced Antibacterial Activity. ChemBioChem 2016, 17 (6), 466–470. https://doi.org/10.1002/cbic.201500674.

(9) Ii, N. F.; Syrb, H.; Vaillancourt, F. H.; Vosburg, D. Dichlorination and Bromination of a Threonyl-S -Carrier Protein by The. ChemBioChem 2006, 7, 748–752. https://doi.org/10.1002/cbic.200500480.

(10) Büchler, J.; Malca, S. H.; Patsch, D.; Voss, M.; Turner, N. J.; Bornscheuer, U. T.; Allemann, O.; Chapelain, C. Le; Lumbroso, A.; Loiseleur, O.; et al. Algorithm-Aided Engineering of Aliphatic Halogenase WelO5* for the Asymmetric Late-Stage Functionalization of Soraphens. Nat. Commun. 2022, 13 (371), 1–11. https://doi.org/10.1038/s41467-022-27999-1.

(11) Matthews, M. L.; Chang, W. C.; Layne, A. P.; Miles, L. A.; Krebs, C.; Bollinger, J. M. Direct Nitration and Azidation of Aliphatic Carbons by an Iron-Dependent Halogenase. Nat. Chem. Biol. 2014, 10 (3), 209–215. https://doi.org/10.1038/nchembio.1438.

(12) Kim, C. Y.; Mitchell, A. J.; Glinkerman, C. M.; Li, F.; Puskal, T.; Weng, J.-K. The Chloroalkaloid (−)-Acutumine Is Biosynthesized via a Fe(II)- and 2- Oxoglutarate-Dependent Halogenase in Menispermaceae Plants. Nat. Commun. 2020, 11 (1867).

(13) Hillwig, M. L.; Liu, X. A New Family of Iron-Dependent Halogenases Acts on Freestanding Substrates. Nat. Chem. Biol. 2014, 10 (11), 921– 923. ttps://doi.org/10.1038/nchembio.1625.

(14) Zhang, Y.; Zhao, C.; Yan, S.; Li, Q.; Zhu, H.; Zhong, Z.; Ye, Y.; Deng, Z. An Iron(II)- and Α-ketoglutarate-dependent Halogenase Acts on Nucleotide Substrates. Angew. Chemie Int. Ed. 2020. https://doi.org/10.1002/anie.201914994.

(15) Zhai, G.; Gong, R.; Lin, Y.; Zhang, M.; Li, J.; Deng, Z.; Sun, J.; Chen, W.; Zhang, Z. Structural Insight into the Catalytic Mechanism of Non-Heme Iron Halogenase AdaV in 2′-Chloropentostatin Biosynthesis. ACS Catal. 2022, 12 (22), 13910–13920. https://doi.org/10.1021/acscatal.2c04608.

(16) Papadopoulou, A.; Meierhofer, J.; Meyer, F.; Hayashi, T.; Schneider, S.; Sager, E.; Buller, R. Re-Programming and Optimization of a L-Proline Cis-4-Hydroxylase for the Cis-3-Halogenation of Its Native Substrate. ChemCatChem 2021, 1–7. https://doi.org/10.1002/cctc.202100591.

(17) Mitchell, A. J.; Dunham, N. P.; Bergman, J. A.; Wang, B.; Zhu, Q.; Chang, W. C.; Liu, X.; Boal, A. K. Structure-Guided Reprogramming of a Hydroxylase to Halogenate Its Small Molecule Substrate. Biochemistry 2017, 56 (3), 441–444. https://doi.org/10.1021/acs.biochem.6b01173.

(18) Neugebauer, M. E.; Kissman, E. N.; Marchand, J. A.; Pelton, J. G.; Sambold, N. A.; Millar, D. C.; Chang, M. C. Y. Reaction Pathway Engineering Converts a Radical Hydroxylase into a Halogenase. Nat. Chem. Biol. https://doi.org/10.1038/s41589-021-00944-x.

(19) Hangasky, J. A.; Taabazuing, C. Y.; Valliere, M. A.; Knapp, M. J. Imposing Function down a (Cupin)-Barrel: Secondary Structure and Metal Stereochemistry in the ΑkG-Dependent Oxygenases. Metallomics 2013, 5 (4), 287–301. https://doi.org/10.1039/c3mt20153h.

(20) Blasiak, L. C.; Vaillancourt, F. H.; Walsh, C. T.; Drennan, C. L. Crystal Structure of the Non-Haem Iron Halogenase SyrB2 in Syringomycin Biosynthesis. Nature 2006, 440 (7082), 368–371. https://doi.org/10.1038/nature04544.

(21) Gorres, K. L.; Pua, K. H.; Raines, R. T. Stringency of the 2-His-1-Asp Active-Site Motifin Prolyl 4-Hydroxylase. PLoS One 2009, 4 (11), 1–6. https://doi.org/10.1371/journal.pone.0007635.

(22) Grzyska, P. K.; Müller, T. A.; Campbell, M. G.; Hausinger, R. P. Metal Ligand Substitution and Evidence for Quinone Formation in Taurine/α-Ketoglutarate Dioxygenase. J. Inorg. Biochem. 2007, 101 (5), 797–808. https://doi.org/10.1016/j.jinorgbio.2007.01.011.

(23) Solomon, E. I.; Deweese, D. E.; Babicz, T. Mechanisms of O2 Activation by Mononuclear Non-Heme Iron Enzymes. Biochemistry 2021, 60, 3497–3506. https://doi.org/10.1021/acs.biochem.1c00370.

(24) Pratter, S. M.; Light, K. M.; Solomon, E. I.; Straganz, G. D. The Role of Chloride in the Mechanism of O 2 Activation at the Mononuclear Nonheme Fe(II) Center of the Halogenase HctB. J. Am. Chem. Soc. 2014, 136, 9385–9395.

(25) Matthews, M. L.; Krest, C. M.; Barr, E. W.; Vaillancourt, H.; Walsh, C. T.; Green, M. T.; Krebs, C.; Bollinger, J. M. Substrate-Triggered Formation and Remarkable Stability of the C - H Bond-Cleaving Chloroferryl Intermediate in the Aliphatic Halogenase, SyrB2. Biochemistry 2009, 48, 4331–4343. https://doi.org/10.1021/bi900109z.

(26) Solomon, E. I.; Goudarzi, S.; Sutherlin, K. D. O2 Activation by Non-Heme Iron Enzymes. Biochemistry 2016, 55 (46), 6363–6374. https://doi.org/10.1021/acs.biochem.6b00635.

(27) Light, K. M.; Hangasky, J. A.; Knapp, M. J.; Solomon, E. I. Spectroscopic Studies of the Mononuclear Non-Heme Fe II Enzyme FIH : Second-Sphere Contributions to Reactivity. J. Am. Chem. Soc. 2013, 135, 9665–9674.

(28) Neidig, M. L.; Brown, C. D.; Light, K. M.; Fujimori, D. G.; Nolan, E. M.; Price, J. C.; Barr, E. W.; Bollinger, J. M.; Krebs, C.; Walsh, C. T.; et al. CD and MCD of CytC3 and Taurine Dioxygenase: Role of the Facial Triad in α-KG-Dependent Oxygenases. J. Am. Chem. Soc. 2007, 129 (46), 14224–14231. https://doi.org/10.1021/ja074557r.

(29) Iyer, S. R.; Chaplin, V. D.; Knapp, M. J.; Solomon, E. I. O2 Activation by Nonheme FeII Α-Ketoglutarate-Dependent Enzyme Variants: Elucidating the Role of the Facial Triad Carboxylate in FIH. J. Am. Chem. Soc. 2018, 140, 11777−11783. https://doi.org/10.1021/jacs.8b07277.

(30) Kal, S.; Que, L. Dioxygen Activation by Nonheme Iron Enzymes with the 2-His-1-Carboxylate Facial Triad That Generate High-Valent Oxoiron Oxidants. J. Biol. Inorg. Chem. 2017, 22 (2–3), 339–365. ttps://doi.org/10.1007/s00775-016-1431-2.

(31) Matthews, M. L.; Neumann, C. S.; Miles, L. A.; Grove, T. L.; Booker, S. J.; Krebs, C.; Walsh, C. T.; Bollinger, J. M. Substrate Positioning Controls the Partition between Halogenation and Hydroxylation in the Aliphatic Halogenase, SyrB2. Proc. Natl. Acad. Sci. U. S. A. 2009, 106 (42), 17723–17728. https://doi.org/10.1073/pnas.0909649106.

(32) Martinie, R. J.; Livada, J.; Chang, W.; Green, M. T.; Krebs, C.; Bollinger, J. M.; Silakov, A. Experimental Correlation of Substrate Position with Reaction Outcome in the Aliphatic Halogenase, SyrB2. J. Am. Chem. Soc. 2015, 137, 6912–6919. https://doi.org/10.1021/jacs.5b03370.

(33) Ryle, M. J.; Padmakumar, R.; Hausinger, R. P. Stopped-Flow Kinetic Analysis of Escherichia Coli Taurine/α-Ketoglutarate Dioxygenase: Interactions with α Ketoglutarate, Taurine, and Oxygen. Biochemistry 1999, 38 (46), 15278–15286. https://doi.org/10.1021/bi9912746.

(34) Price, J. C.; Barr, E. W.; Tirupati, B.; Bollinger, J. M.; Krebs, C. The First Direct Characterization of a High-Valent Iron Intermediate in the Reaction of an α-Ketoglutarate-Dependent Dioxygenase: A High-Spin Fe(IV) Complex in Taurine/α-Ketoglutarate Dioxygenase (TauD) from Escherichia Coli. Biochemistry 2003, 42 (24), 7497–7508. https://doi.org/10.1021/bi030011f.

(35) Ho, R. Y. N.; Mehn, M. P.; Hegg, E. L.; Liu, A.; Ryle, M. J.; Hausinger, R. P.; Que, L. Resonance Raman Studies of the Iron(II)-α-Keto Acid Chromophore in Model and Enzyme Complexes. J. Am. Chem. Soc. 2001, 123 (21), 5022–5029. https://doi.org/10.1021/ja0041775.

(36) Saban, E.; Chen, Y. H.; Hangasky, J. A.; Taabazuing, C. Y.; Holmes, B. E.; Knapp, M. J. The Second Coordination Sphere of FIH Controls Hydroxylation. Biochemistry 2011, 50 (21), 4733–4740. https://doi.org/10.1021/bi102042t.

(37) Chaplin, V. D.; Hangasky, J. A.; Huang, H. T.; Duan, R.; Maroney, M. J.; Knapp, M. J. Chloride Supports O2 Activation in the D201G Facial Triad Variant of Factor-Inhibiting Hypoxia Inducible Factor, an α-Ketoglutarate Dependent Oxygenase. Inorg. Chem. 2018, 57 (20), 12588–12595. https://doi.org/10.1021/acs.inorgchem.8b01736.

(38) Mondal, P.; Rajapakse, S.; Wijeratne, G. B. Following Nature’s Footprint: Mimicking the High-Valent Heme-Oxo Mediated Indole Monooxygenation Reaction Landscape of Heme Enzymes. J. Am. Chem. Soc. 2022, 144 (9), 3843–3854. https://doi.org/10.1021/jacs.1c11068.

(39) Caranto, J. D.; Weitz, A.; Hendrich, M. P.; Kurtz, D. M. The Nitric Oxide Reductase Mechanism of a Flavo-Diiron Protein: Identification of Active-Site Intermediates and Products. J. Am. Chem. Soc. 2014, 136 (22), 7981–7992. https://doi.org/10.1021/ja5022443.

(40) Hosseinzadeh, P.; Marshall, N. M.; Chacón, K. N.; Yu, Y.; Nilges, M. J.; New, S. Y.; Tashkov, S. A.; Blackburn, N. J.; Lu, Y. Design of a Single Protein That Spans the Entire 2-V Range of Physiological Redox Potentials. Proc. Natl. Acad. Sci. U. S. A. 2016, 113 (2), 262–267. https://doi.org/10.1073/pnas.1515897112.

(41) Crameri, F.; Shephard, G. E.; Heron, P. J. The Misuse of Colour in Science Communication. Nat. Commun. 2020, 11 (1), 1–10. https://doi.org/10.1038/s41467-020-19160-7.

