## Supplementary Information for "Electrostatically regulated active site assembly governs reactivity in non-heme iron halogenases"

### Table of Contents

**Experimental Procedures**……………………………………………………………………….S2

**1. Protein expression and purification**……………………………………..…………….S2

**2. Anaerobic Equilibrium UV-Vis studies**……………………………….……………..S3

**3. Equilibrium Dialysis Studies**……………………………….……………….……..S3

**4. Stopped-flow UV-Vis spectroscopy**…………………………….………………..…S4

**5. Molecular dynamics**……………………………..……………….…………………S4

**6. DFT calculations**…………………………………………………………....……………S5

**7. Anion activity assays**……………………………………………….……..……….….S6

**8. Accurate Mass Determination of BesD Enzymatic Products**……………………..…….S6

**Supplementary Tables**……………………………………………………………………S9

**Table S1.** SRM transitions monitored for UPLC-MS/MS detection of Lysine and enzymatically derivatized Lysine ……..…………………………………………………………..……………S9

**Table S2.** Accurate Mass Determination of BesD Enzymatic Product……..……………….S10

**Supplementary Figures**……………..…………………………………………………..…S11

**Figure S1.** Substrate-anion cooperativity in HalB……………..…………..………………..….S11

**Figure S2.** AQC-derivatized lysine HPLC standard curve………..………..………………..….S12

**Figure S3**. Equilibrium dialysis HPLC traces.……………..…...…...……....………………..….S13

**Figure S4.** Chloride titration MLCT band spectra…………………….…………………….S14

**Figure S5.** Electrostatic surfaces generated from DFT optimized MD simulation snapshots……………………………………………………………………………………..S15

**Figure S6.** UPLC-MRM-MS/MS chromatograms of brominated lysine and other analytes..S16

**Figure S7.** UPLC-MRM-MS/MS chromatograms of azidated lysine and other analytes …S17

**Figure S8.** UPLC-MRM-MS/MS chromatograms of nitrated lysine and other analytes…...S18

**References**……………..…………………………………………………….……………S19

### Experimental Procedures

#### 1. Protein expression and purification

Plasmid for His10-tagged *Streptomyces lavenduligriseus* BesD and *Streptomyces wuyuanensis* HalB contained in a pET16b vector was received as a gift from Chang lab (UC Berkeley) and sequenced to confirm gene integrity. The plasmids were transformed into Rosetta 2 (DE3) cells for protein overexpression. Colonies from an overnight LB-agar plate containing carbenicillin (50 ug $ml^{-1}$) and chloramphenicol (37 ug $ml^{-1}$) were inoculated into 250 ml flasks containing TB media (50 ml) with the same concentrations of antibiotics and incubated at 37˚ C overnight at 200 RPM. Culture flasks with TB (1.5 L) and appropriate antibiotics were inoculated with overnight cultures and grown at 37˚ C at 200 RPM until cells reached an $OD_{600}$ of 0.9 - 1.1. Upon reaching appropriate optical density, cultures were cooled on ice for 15 minutes before the addition of IPTG (0.25 mM) and then incubated at 16˚ C for 24 hours and 18 hours for BesD and HalB respectively. After overexpression, cells were collected by centrifugation at 8000 RPM at 4˚ C, flash frozen, and stored at -20˚ C for further use.

Frozen cells were re-suspended in lysis buffers (50 mM HEPES, 300 mM NaCl, 10 mM imidazole, 1 mM $CaCl_2$, 25 mM $MgCl_2$, 0.1 mg/ml RNase, and 0.1 mg/ml Dnase, pH = 7.5) and then lysed by sonication. Lysed cell solution was centrifuged to separate out insoluble cell debris, and then supernatant was collected and filtered through 0.45-micron filters. Filtered supernatant was applied to a 5 mL HisTrapFF column using an AKTA Start protein purification system. After sample application, column was washed with 15 column volumes (CVs) of wash buffer (50 mM HEPES, 300 mM NaCl, 10 mM imidazole, and 20 mM 2-mercaptoethanol, pH = 7.5), and then His-tagged protein was eluted using a two-step gradient method first with 45% elution buffer (50 mM HEPES, 300 mM NaCl, 300 mM imidazole, and 20 mM 2-mercaptoethanol, pH = 7.5) for 8 CVs to remove non-specific interacting proteins which was then increased to 100% elution buffer to elute high purity protein for 8 CVs. Fractions from the last elution step were collected and dialyzed in 50 mM HEPES, 100 mM NaCl, 1 mM DTT, and 1 mM EDTA (pH = 7.5) for 2 hours and then transferred to a dialysis buffer of 50 mM HEPES, 1 mM DTT, 100 mM NaCl (pH = 7.5) for 2 hours. After this second round of dialysis, protein solution was diluted to a concentration of 0.05 mM and then incubated with HRV-3C protease (1 mg protease to 100 mg protein) dialyzing against 50 mM HEPES, 1 mM DTT, 100 mM NaCl (pH = 7.5) overnight. Imidazole was added to the protein solution so that the final concentration would be 10 mM. Protein solution was applied to a 5 mL HisTrapFF column using an AKTA Start protein purification system. His-tagged cleaved BesD was recovered during sample application and subsequent 4 CV wash with wash buffer.

Protease and residual His-tagged protein were eluted with elution buffer. Collected fractions were concentrated using Amicon Ultra-15 Centrifugal Filter (10 kDa MWCO). Concentrated protein was applied to a PD-10 desalting column equilibrated with 50 mM HEPES (pH = 7.5) and fractions were collected based on absorbance at 280 nm. Collected fractions were further concentrated, and glycerol (15%) was added to the protein solution for storage. BesD and HalB's purity and His-tag cleavage was confirmed via gel electrophoresis and mass spectrometry. Protein solution was aliquoted, flash frozen, and stored at -80˚ C for further use.

**2. Anaerobic Equilibrium UV-Vis studies**

All UV-Vis studies were performed anaerobically on a Cary60 UV-Vis spectrophotometer at 11˚ C. Anaerobic solutions of 100 mM HEPES (pH = 7.5) and 100 mM 2OG in 100 mM HEPES (pH=7.5) were prepared through alternating cycles of vacuum degassing followed by argon flushes. Dry solids for all other compounds were brought into the anaerobic glovebag environment and solutions were prepared with the $O_2$-free solution stocks.

Before each UV-Vis experiment, BesD or HalB was buffer exchanged three times into $O_2$-free 100 mM HEPES (pH = 7.5) in 0.5 mL Amicon™ Ultra 10 kDa MWCO centrifugal filter units by centrifugation at 13500 RPM for 20 minutes at 4˚ C. A solution of BesD or HalB was prepared in a cuvette to a final concentration of 0.13 mM ± 0.02. Aliquots of $(NH_4)_2Fe(SO_4)_2$ in water, 2-oxoglutarate in 100 mM HEPES, NaX (X = Cl, Br, $N_3$, $NO_2$) in 100 mM HEPES, and L-lysine in 100 mM HEPES were added sequentially to the protein solution to final concentrations of 97.5 µM $(NH_4)_2Fe(SO_4)_2$, 630 µM 2OG, 5 mM NaX (X = Cl, Br, $N_3$, $NO_2$), and 5 mM L-lysine. Stock solutions of each compound were prepared so that small aliquots could be added to the cuvette and keep the volume change below 10%. Multiple spectra were acquired 10 minutes after each component addition to ensure stable analyte signal. All spectra shown are background corrected by the average signal of 775 nm to 800 nm and dilution corrected unless otherwise stated.

Titrations were performed analogous to UV-Vis addition experiments but final concentrations in the cuvette before the additions of chloride titrant solutions were 330 µM BesD, 600 µM $(NH_4)_2Fe(SO_4)_2$, and 800 µM 2OG in 100 mM HEPES (pH = 7.5) at lysine concentrations indicated in figures. Increases in absorbance at 518 nm were converted into fraction of chloro-bound protein by dividing the titrant absorbance value by the theoretical max, and the resulting scaled data was fit using the quadratic binding equation.

**3. Equilibrium Dialysis Studies**

All equilibrium dialysis experiments were conducted anaerobically. To prepare the protein, BesD was brought into a glovebag and buffer exchanged three times into $O_2$-free 50 mM HEPES (pH = 7.5) as described previously. Stock solutions of compounds were prepared as described in the previous section. Protein solutions of 400-450 μM BesD, 4 mM 2OG, 1 mM $(NH_4)_2Fe(SO_4)_2$, 0 or 30 mM NaCl, and 45 mM or 35 mM $Na_2SO_4$ in 50 mM HEPES (pH = 7.5) were dialyzed against an equal volume of lysine containing solution (125 μM – 1000 μM) in 4 mM 2OG, 1 mM $(NH_4)_2Fe(SO_4)_2$, 0 or 30 mM NaCl, 45 mM or 35 mM $Na_2SO_4$, and 50 mM HEPES (pH = 7.5) for 20 hours. Concentrations of NaCl and $Na_2SO_4$ depended on the identified experiment as detailed in **Fig S3**. After 20 hours, 50 μL of samples from the non-protein chamber were combined with 40 μL of 30 mM borate buffer (pH = 10.5) and then amino acids were derivatized with 10 μL of ca. 10 mM 6-aminoquinolyl-N-hydroxysuccinimidylcarbamate (AQC) in acetonitrile solution. Immediately after derivatizing agent was added, samples were vortexed for 10 seconds. Samples with initial lysine concentrations 800-1000 μM were diluted 2-fold, and 50 μL from the diluted sample was derivatized and analyzed.

Derivatized amino acid samples were then run on a Shimadzu Prominence-i LC-2030C 3D Plus system equipped with a Regis Technologies REXCHROM C18 column (4.60 mm × 250 mm × 5 μm) and a PDA detector. Separations occurred with a gradient separation at a temperature of 35 °C, an injection volume of 25 or 50 μL (specific injection volumes are indicated in relevant figure captions), and a flow rate of 0.5 mL/min. The separation method is as follows: 0% B to 70% B, 0.0 to 40.0 min; 70% B to 100% B, 40.0 to 42.0 min; 100% B, 42.0 to 50.0 min; 100% B to 0% B, 50.0 to 52.0 min; 0% B, 52.0 to 60.0 min, where solution A is 5 mM ammonium acetate and solution B is 60% acetonitrile in water. All HPLC traces were constructed from absorbance at 254 nm. Peak areas were converted to concentrations through a standard curve (**Fig. S2**) to determine the free lysine concentration in each equilibrium dialysis sample, allowing the data to be fit by the following Hills's equation model:

$$\frac{[ES]}{[E]_t} = \frac{[S]}{[S] + K_{d,app}} \tag{1}$$

Where [S] is the free ligand concentration, [ES] is the ligand-protein complex concentration, $[E]_t$ is the total enzyme concentration, and $K_{d,app}$ is the apparent binding affinity.

#### 4. Stopped-flow UV-Vis spectroscopy

Protein solutions for stopped-flow studies prepared similarly to equilibrium studies. Before stopped-flow mixing, final solutions of anaerobic protein solutions were 225 μM BesD, 2.25 mM Lysine, 10 mM NaCl, 170 μM $(NH_4)_2Fe(SO_4)_2$, 2.25 mM 2OG in 100 mM HEPES (pH = 7.5) unless the experiment indicates a component was absent during the measurement. Room temperature $O_2$ saturated solutions of 100 mM HEPES (~1 mM) were prepared by bubbling $O_2$ gas through solution for several hours. Stopped-flow was run on an Applied Photophysics SX-20 paired with an Ocean Optics detector triggered through a Siglent waveform generator. Equal volumes of prepared anaerobic protein solution and $O_2$-saturated buffer were combined at 11˚C in an anaerobic glovebag.

### 5. Molecular dynamics

Homology model for BesD from *Streptomyces lavenduligriseus* was created using the SWISS-MODEL tool[1] with default parameters from the crystal structure of BesD from *Streptomyces cattleya* which shares 75% identity with target protein.[1–6] Hydrogen atoms were added to homology model at a pH of 7.5 with the H++ webserver.[7–9] Using the tleap module in AMBER 20, BesD was parameterized with the ff19SB force field, 2OG was parameterized using GAFF (charges were assigned with the antechamber module), and iron was described as a ferrous species.[10–14] The protein was solvated in a 10.0 Å OPC water box, and Na+ and Cl- ions were added to neutralize the system.[15] Substrate lysine was treated as a zwitterionic species with a charged epsilon amine.[16] All varying coordination systems were created manually by editing input pdb files prior to the creation of AMBER input files with the tleap module. The systems representing the varied coordination states were minimized (first solvent, then protein), gently heated to 300 K, and density was equilibrated for 2 ns. For each coordination system, two independent 250 ns production runs were performed. To maintain the primary coordination sphere of iron in the six-coordinate systems, all ligand distances were restrained to within 0.1 Å of their crystallographic distances using a force constant of 100.0 kcal mol-1 Å-1. For MD simulations involving the five-coordinate (5C) species, the iron-ligand bonds and angles were treated explicitly using parameters developed from the MCPB python package in AmberTools21.[17,18] For all MCPB calculations, the B3LYP/6-31G(d) level of theory was employed.[19–21] During geometry optimizations, the alpha carbons of all amino acids and C5 of the alpha-ketoglutarate ligand were held fixed to mimic their positions within the protein structure. Trajectory analyses were performed with CPPTRAJ; H-bond donor-acceptor cutoff distances were set to 3.2 Å.[22] Error bars in the analysis were calculated using the standard error over the independent simulations (standard deviation divided by √2).[23]

### 6. DFT calculations

All electronic structure calculations were performed with the ORCA software package (version 5.0.3).[24,25] All geometry optimizations were conducted at the B3LYP-D3BJ/def2-SVP level of theory; iron and all coordinating atoms in the primary coordination sphere were treated with the def2-TZVP basis set.[19,20,26–29] Revised BJ damping parameters were employed to prevent over-correction of the treatment of dispersion interactions.[30] The resolution of identity approximation for coulomb and numerical chain-of-sphere integration for exchange integrals (keyword: RIJCOSX) were used to accelerate all geometry optimizations. The auxiliary basis def2/J was called for all geometry optimizations.[31] The keyword 'defgrid2' was used to build all integration grids, and the keywords 'SOSCF' and 'SlowConv' were used to aid the SCF convergence.

Representative snapshots from each MD trajectory where E119 was a median distance away from the iron center were taken as the input to build DFT models for different active site configurations. The model systems include iron, all ligands coordinated to it, substrate lysine, and protein residues Asn218 and Glu119. All protein residues were truncated and capped at the alpha carbon. 2OG was truncated to pyruvate to prevent unphysical interactions during geometry optimizations. The alpha carbons of all amino acid residues and the methyl carbon of pyruvate were held fixed during the geometry optimization. The protein environment was simulated using a dielectric constant of $\varepsilon = 4$ using the CPCM solvation model.

After geometry optimization, the Multiwfn package was used to calculate RESP charges through a two-step procedure and then used to generate electron density and electrostatic potential surfaces.[32–34] The resulting surfaces were visualized in VMD with a surface Isovalue of 0.02 and color scale data range set to 0.20 to -0.20.[35]

#### 7. Anion activity assays

All reactions were prepared anaerobically. BesD was brought into a glovebag after six antechamber cycle and allowed to equilibrate in an anaerobic environment for 1 hour. For non-native anion reactions, BesD (final concentration: 50 μM) was combined with 10 mM anion salt (NaBr, $NaN_3$, or $NaNO_2$), 5 mM 2OG, 1 mM sodium ascorbate, 100 μM $(NH_4)_2Fe(SO_4)_2$, and 200 μM lysine. All samples were brought out of the glovebag and exposed to oxygen to initiate reactions. Reactions containing azide and nitrite were incubated in an Eppendorf Thermomixer® at 23˚ C overnight mixing (350 RPM). Due to the observed instability of the brominated species, reactions containing bromide were incubated in an Eppendorf Thermomixer® at 4˚ C for 1 hour mixing (350 RPM).

After incubations, all reactions were filtered through 0.5 mL Amicon™ Ultra 10 kDa MWCO centrifugal filter units at 13500 RPM for 20 minutes at 4˚ C. Reaction flowthrough (135 μL) was collected for further processing from each reaction.

Collected reaction flowthrough samples (135 μL) were combined with 20 μL of 100 mM sodium tetraborate (pH=8.5), 5 μL of 200 μM of L-Lysine(13C6, 99%; 15N2, 99%)·2HCl, 1.5 μL 5 M NaOH, and 18.5 μL of MilliQ $H_2O$. AQC in acetonitrile was added to each reaction to a final concentration of ca. 1 mM and a total sample volume of 200 µL. Samples were vortexed immediately after AQC addition. To analyze reaction product distributions, reaction samples were run on a Waters Acquity UPLC coupled to a Waters triple quadrupole mass spectrometer (Acquity TQD). Column choice, gradient method, and instrument parameters were adapted from Ref 36.[36] Mass transitions monitored for each reaction are tabulated in **Table S1**.

**8. Accurate Mass Determination of BesD Enzymatic Products**

A Sciex Exion UHPLC coupled to a Sciex X500R quadrupole time-of-flight (qtof) mass spectrometer was used for separation and accurate mass measurement of AQC-derivatized BesD enzymatic products. A Waters CORTECS UPLC C18 2.1 mm x 100 mm column (1.6 mm particles) at 55 °C was used during the following 10 min gradient separation with A: Water containing 0.1% formic acid and B: ACN containing 0.1% formic acid, at a flow rate of 0.5 mL/min: 1% B, 0 min to 1.0 min; 1% B to 13% B, 1.0 min to 2.0 min; 13% B to 15% B, 2.0 min to 5.5 min; 15%B to 95% B, 5.5. min to 6.5 min; 95% B, 6.5 min to 7.5 min; 95% B to 1% B, 7.5 min to 7.7 min; 1% B, 7.7 min to 10 min. Electrospray ionization mass spectra in positive ionization mode were collected over the range m/z 50-1200 during the analysis. MS parameters were as follows: Ion source gas 1: 45 psi; Ion source gas 2: 45 psi; Curtain gas: 30 psi; CAD gas: 7; Temp: 500 °C; Spay voltage: 550V; Declustering potential: 50V; DP spread: 0V; CE: 10V; CE spread: 0V.

**Table S1.** SRM transitions monitored for UPLC-MS/MS detection of Lysine and enzymatically derivatized Lysine.

| Compound | $[M+xH]^{x+}$ | SRM Transition | Cone (V) | Collision Energy (eV) |
|---|---|---|---|---|
| AQC(2x)-Lysine | $[M+2H]^{2+}$ | 244.1>171.1 | 35 | 20 |
| AQC(2x)-OH-Lysine | $[M+2H]^{2+}$ | 252.1>171.1 | 35 | 20 |
| AQC(2x)-Lac-Lysine | $[M+2H]^{2+}$ | 243.1>171.1 | 35 | 20 |
| AQC(2x)-35Cl-Lysine | $[M+2H]^{2+}$ | 261.1>171.1 | 35 | 20 |
| AQC(2x)-37Cl-Lysine | $[M+2H]^{2+}$ | 262.1>171.1 | 35 | 20 |
| AQC(2x)-$N_3$-Lysine | $[M+2H]^{2+}$ | 264.6>171.1 | 35 | 20 |
| AQC-$N_3$-Lysine-AQC | $[M+H]^{+}$ | 528.2>171.1 | 35 | 20 |
| AQC(2x)-79Br-Lysine | $[M+2H]^{2+}$ | 283.1>171.1 | 35 | 20 |
| AQC(2x)-81Br-Lysine | $[M+2H]^{2+}$ | 284.1>171.1 | 35 | 20 |
| AQC(2x)-$NO_2$-Lysine | $[M+2H]^{2+}$ | 266.6>171.1 | 35 | 20 |
| AQC-$NO_2$-Lysine-AQC | $[M+H]^{+}$ | 532.2>171.1 | 35 | 20 |

**Table S2.** Accurate Mass Determination of BesD Enzymatic Products

| Analyte | Chemical formula | Theoretical $[M+H]^{1+}$ | Observed $[M+H]^{1+}$ | Mass error (ppm) | Theoretical $[M+2H]^{2+}$ | Observed $[M+2H]^{2+}$ | Mass error (ppm) |
|---|---|---|---|---|---|---|---|
| AQC(2x)-Lac-Lysine | $C_{26}H_{24}N_6O_4$ | 485.1932 | 485.1932 | 0.000 | 243.1003 | 243.0994 | -3.538 |
| AQC(2x)-OH-Lysine | $C_{26}H_{26}N_6O_5$ | 503.2038 | 503.2042 | 0.795 | 252.1056 | 252.1051 | -1.983 |
| AQC(2x)-Cl-Lysine | $C_{26}H_{25}ClN_6O_4$ | 521.1699 | 521.17 | 0.192 | 261.0886 | 261.0879 | -2.681 |
| AQC(2x)-Br-Lysine | $C_{26}H_{25}BrN_6O_4$ | 565.1194 | 565.1204 | 1.770 | 283.0634 | 283.0633 | -0.353 |
| AQC(2x)-$N_3$-Lysine | $C_{26}H_{25}N_9O_4$ | 528.2103 | 528.2099 | -0.757 | 264.6088 | 264.6082 | -2.267 |

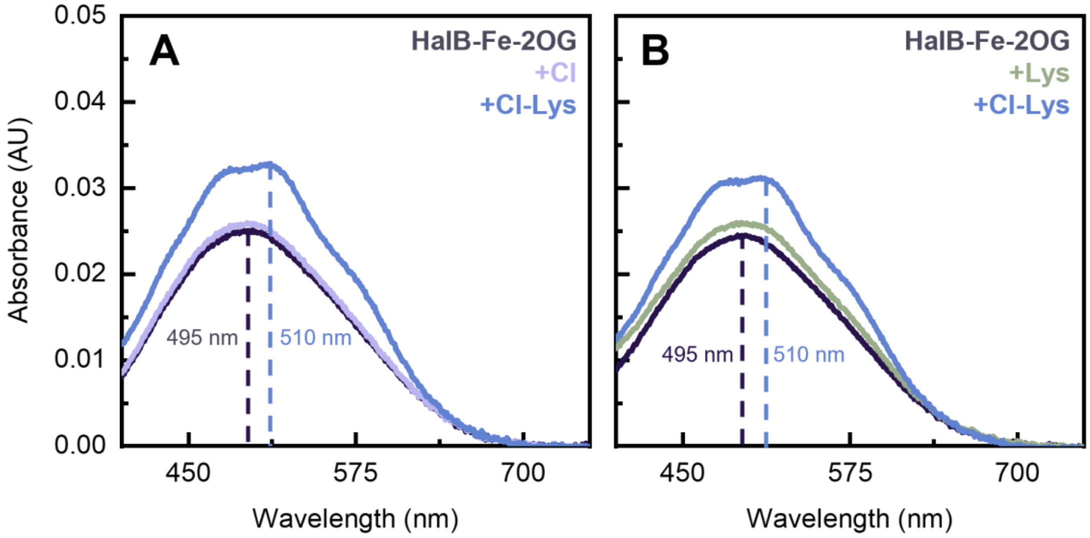


**Figure S1.** Substrate-anion cooperativity in HalB. Spectral changes in MLCT band of 130±5 μM HalB, 97.5 μM Fe, 630 μM 2OG in 100 mM HEPES (pH =7.5) upon sequential additions of **(A)** 5 mM Cl then 5 mM lysine or **(B)** 5 mM lysine then 5 mM Cl. Lack of a perturbation in the MLCT band until the addition of Lys/Cl demonstrates that substrate-anion cooperativity is operant in HalB as well as BesD. Slight upward shift upon addition of 5mM lysine in **(B)** is due to intrinsic absorbance of lysine in this region.

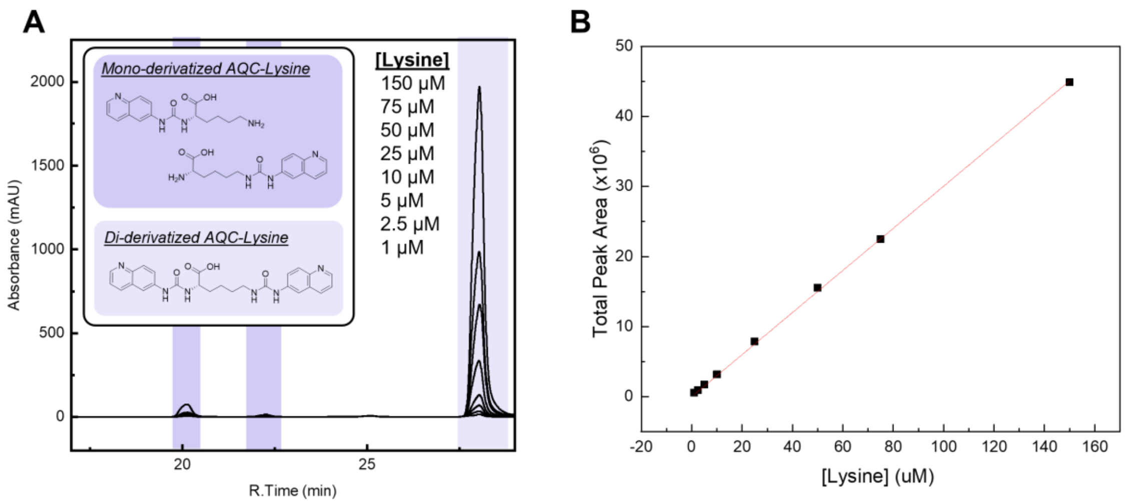


**Figure S2.** AQC-derivatized lysine HPLC standard curve. **(A)** HPLC chromatograms of standard lysine samples (1-150 µM). AQC reacts with amine functional groups and therefore lysine derivatization with AQC gives three products as shown. Peak area integrations were performed in Shimadzu lab solution software. Injection volume was 50 µls. **(B)** Total adjusted peak area of all three AQC-tagged lysine peaks graphed versus concentration of lysine. Mono-derivatized AQC-lysine products were approximated to have half the absorbance of the di-derivatized AQC-lysine product and therefore peak area corresponding to the monoderivative species were multiplied by 2 when summed to create calibration curve.

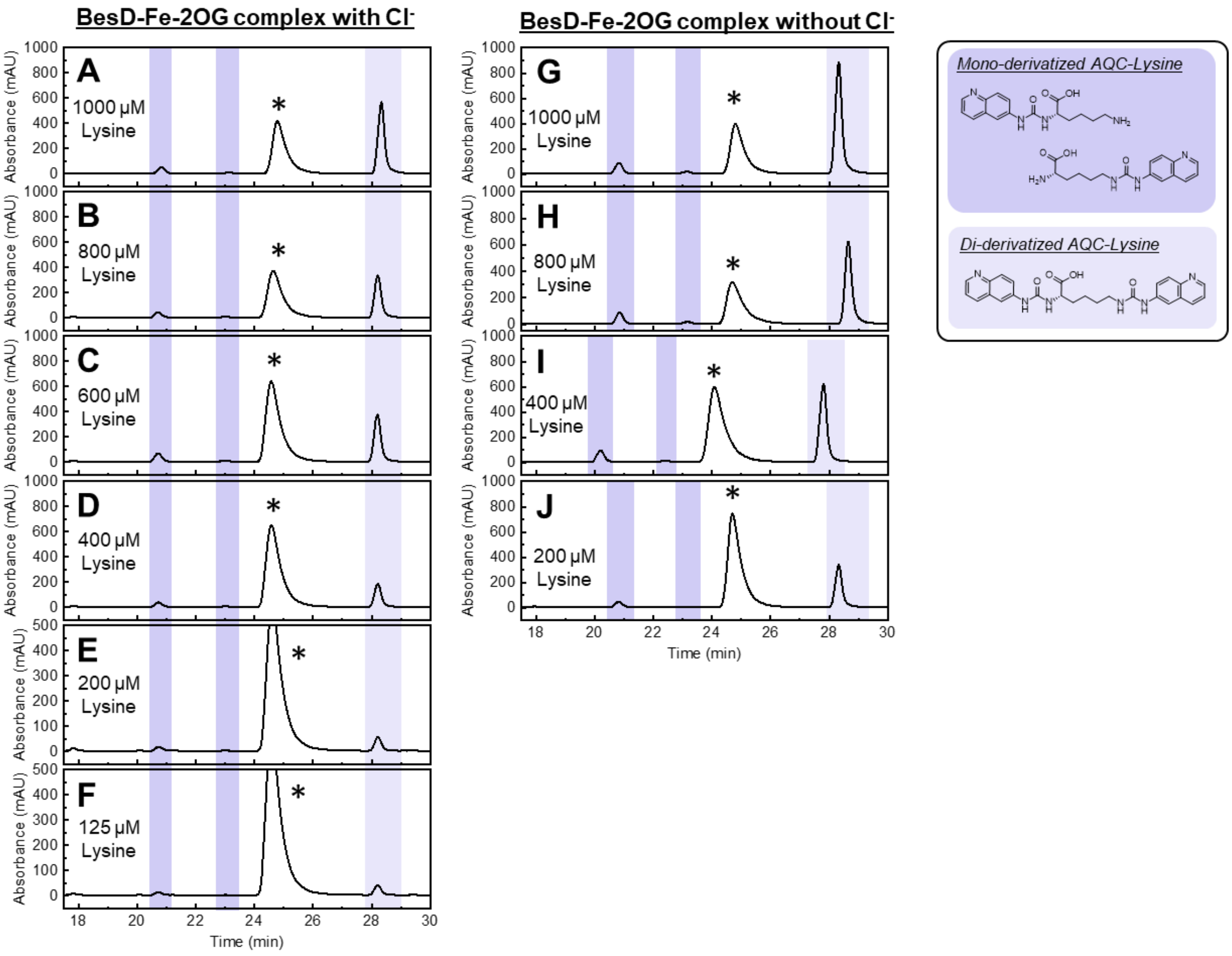


**Figure S3.** Equilibrium dialysis HPLC chromatograms. HPLC chromatograms of samples from **(A-F)** chloride containing equilibrium dialysis condition (4 mM 2OG, 1 mM $(NH_4)_2Fe(SO_4)_2$, 30 mM NaCl, 35 mM $Na_2SO_4$ in 50 mM HEPES (pH = 7.5) ) and from **(G-J)** chloride-absent equilibrium dialysis condition (4 mM 2OG, 1 mM $(NH_4)_2Fe(SO_4)_2$, 45 mM $Na_2SO_4$ in 50 mM HEPES (pH = 7.5). All samples were taken from the protein-free side of the equilibrium dialysis chamber and then derivatized with AQC. Injection volume onto the HPLC was 25 µls. Initial lysine concentrations before equilibrium are provided as an inset in the chromatogram. Peak area for all three possible AQC-lysine products were combined by a weighted summation to account for the difference in absorbance between the di-derivatized lysine and the mono-derivatized lysine product as describe in S2. The weighted summation of the three peak areas of AQC-lysine products was converted to concentration using the AQC-lysine calibration curve, adjusted for 25 µl injections. Peaks identified with an asterisks are products of the AQC-tagging reaction with ammonium from $(NH_4)_2Fe(SO_4)_2$. Chromatogram I was obtained without a guard column which is why the retention times for peaks deviate from the rest of the samples.

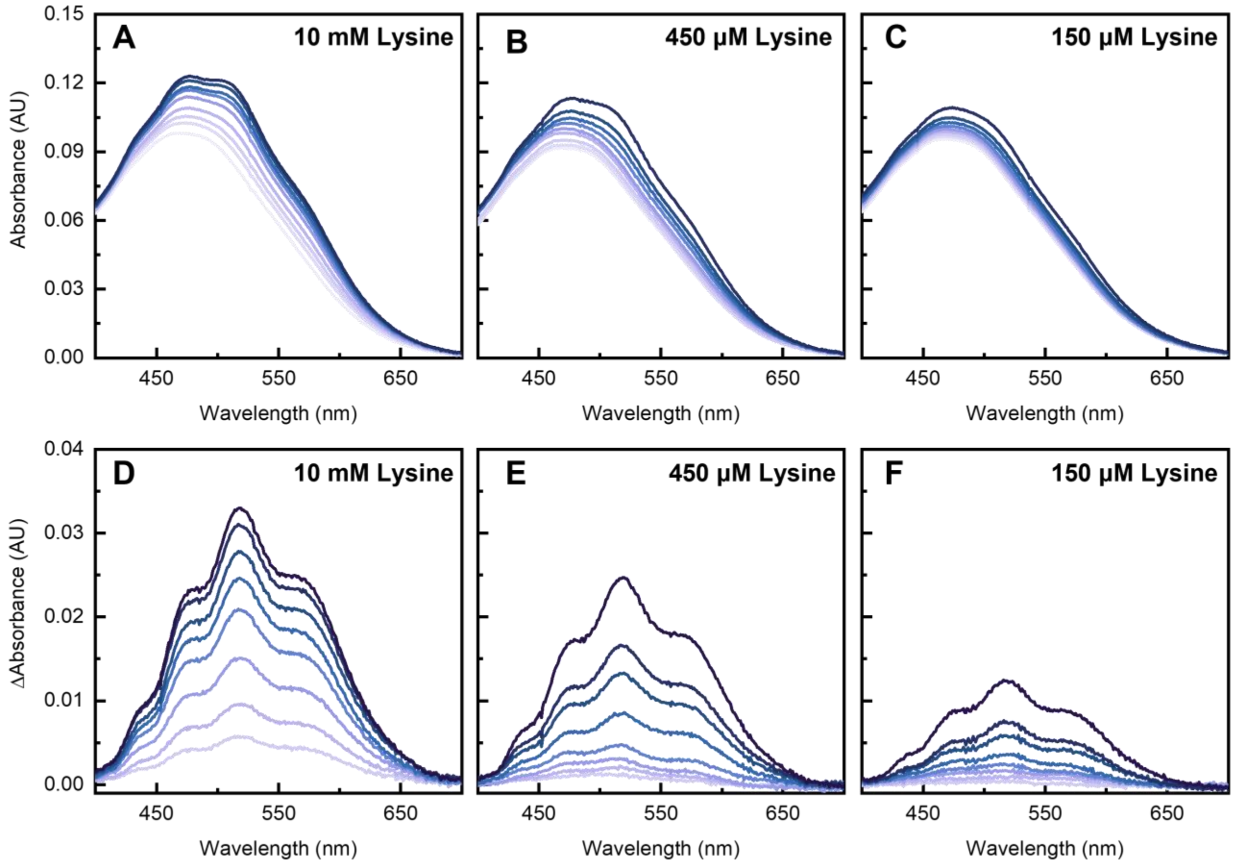


**Figure S4.** Chloride titration MLCT band spectra. **(A-C)** Background and dilution corrected spectra of chloride titrations (final concentration in cuvette of 0-25 mM) into 330 μM BesD, 600 μM $(NH_4)_2Fe(SO_4)_2$, and 800 μM 2OG in 100 mM HEPES (pH = 7.5) at different concentrations of lysine (0.15-10 mM). **(D-F)** Difference spectra of titration data where initial 0 mM Cl spectra has been subtracted from subsequent chloride additions spectra. Spectra have been background corrected to a line defined by absorbances at 390 nm and 750 nm. Increases in absorbance at 518 nm from the difference spectra were used to construct the titration curve for chloride binding at various lysine concentrations.

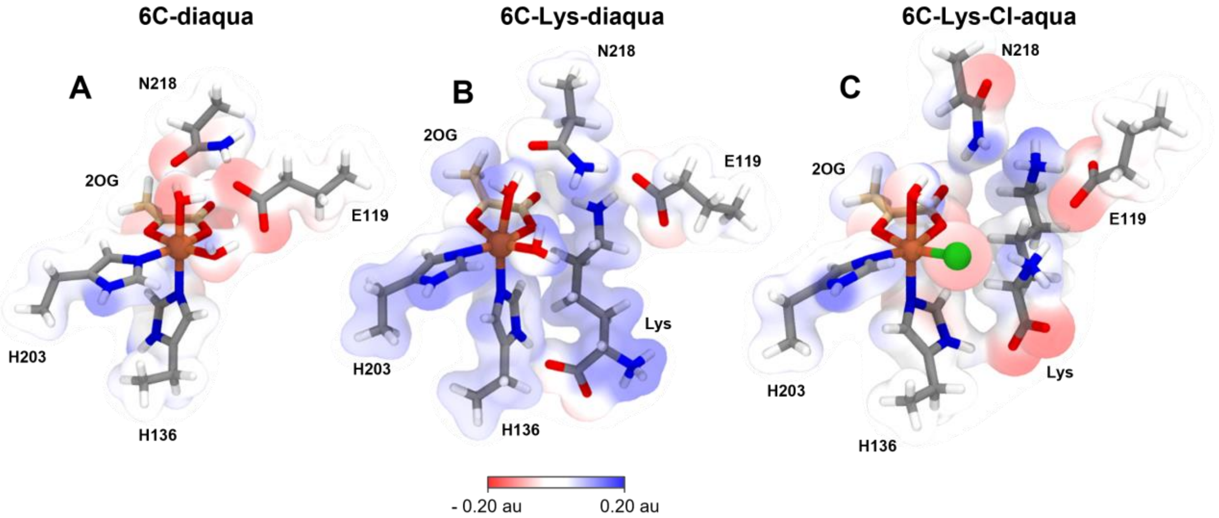


**Figure S5.** Electrostatic surfaces of BesD active site configurations. Electrostatic surfaces for **(A)** the *6c*-diaqua species, **(B)** the *6c*-diaqua species in the presence of lysine, and **(C)** the *6c*-Cl-aqua species in the presence of lysine were generated from DFT optimized of representative MD simulation snapshots. The aqua ligand in **(C)** was added back in before the geometry optimization for more consistent comparisons between surfaces. N218 was found to form frequent hydrogen bond interactions with the axial aqua ligand and was included as it is known to be an important residue in this enzyme in addition to the MD considerations. After geometry optimization, the Multiwfn package was used to calculate RESP charges and generate electron density/electrostatic potential surfaces. Surfaces were visualized in VMD with a surface Isovalue of 0.02 and color scale data range set to 0.20 to -0.20.

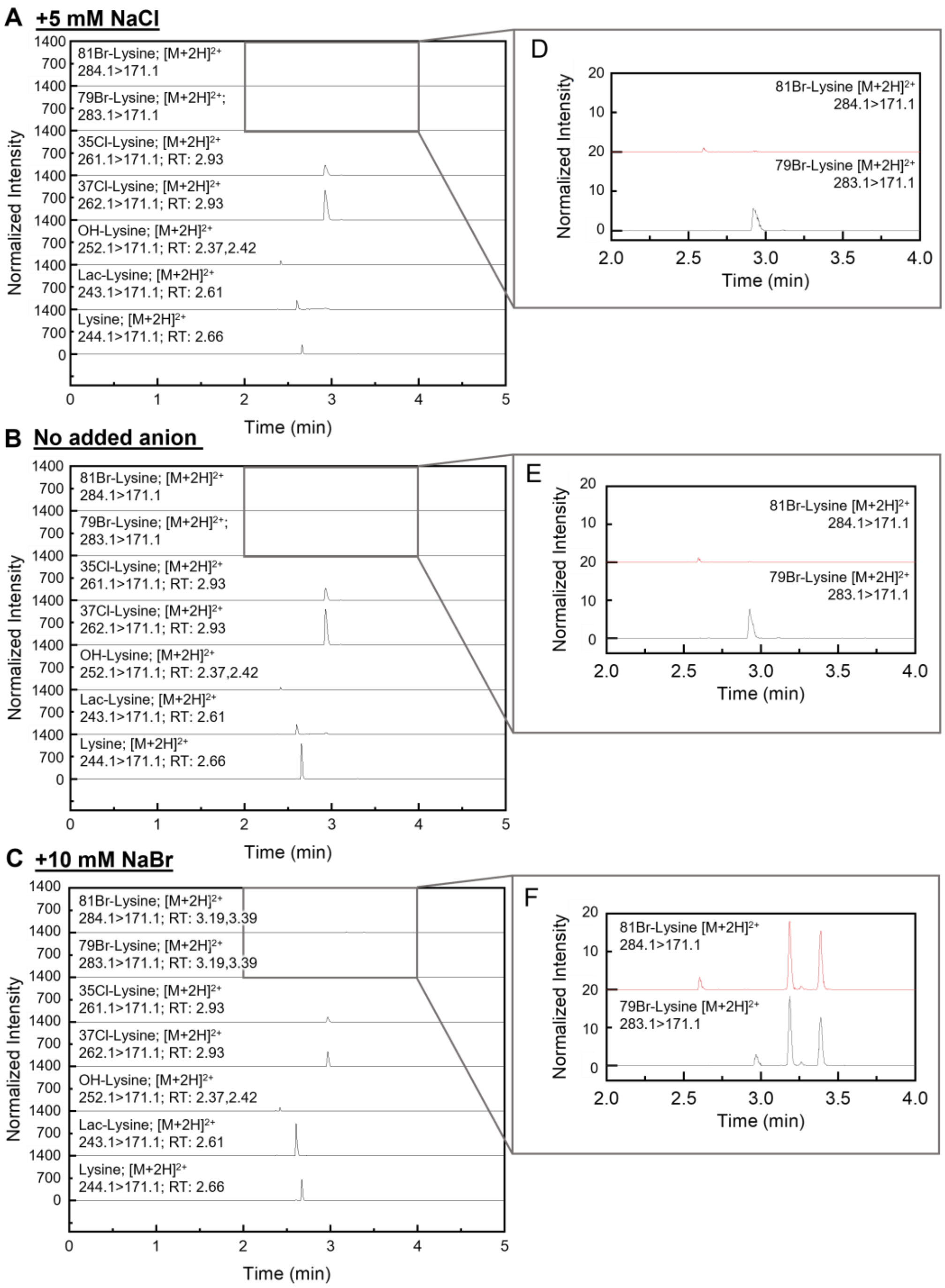


**Figure S6.** UPLC-MRM-MS/MS chromatograms of brominated lysine and other analytes. Reaction samples were prepared with 50 μM BesD, 5 mM 2OG, 1 mM sodium ascorbate, 100 μM $(NH_4)_2Fe(SO_4)_2$, 200 μM lysine in 100 mM HEPES (pH = 7.5) with either **(A)** 5 mM NaCl, **(B)** no added exogenous anion, or **(C)** 10 mM NaBr. Zoomed chromatograms of brominated lysine product channels **(D-F)** show peaks for brominated products only appear when bromide is directly added to the reaction. Accurate masses are confirmed in **Table S2**. All chromatograms shown have had intensity normalized to internal standard signal (L-Lysine(13C6, 99%; 15N2, 99%)·2HCl) intensity.

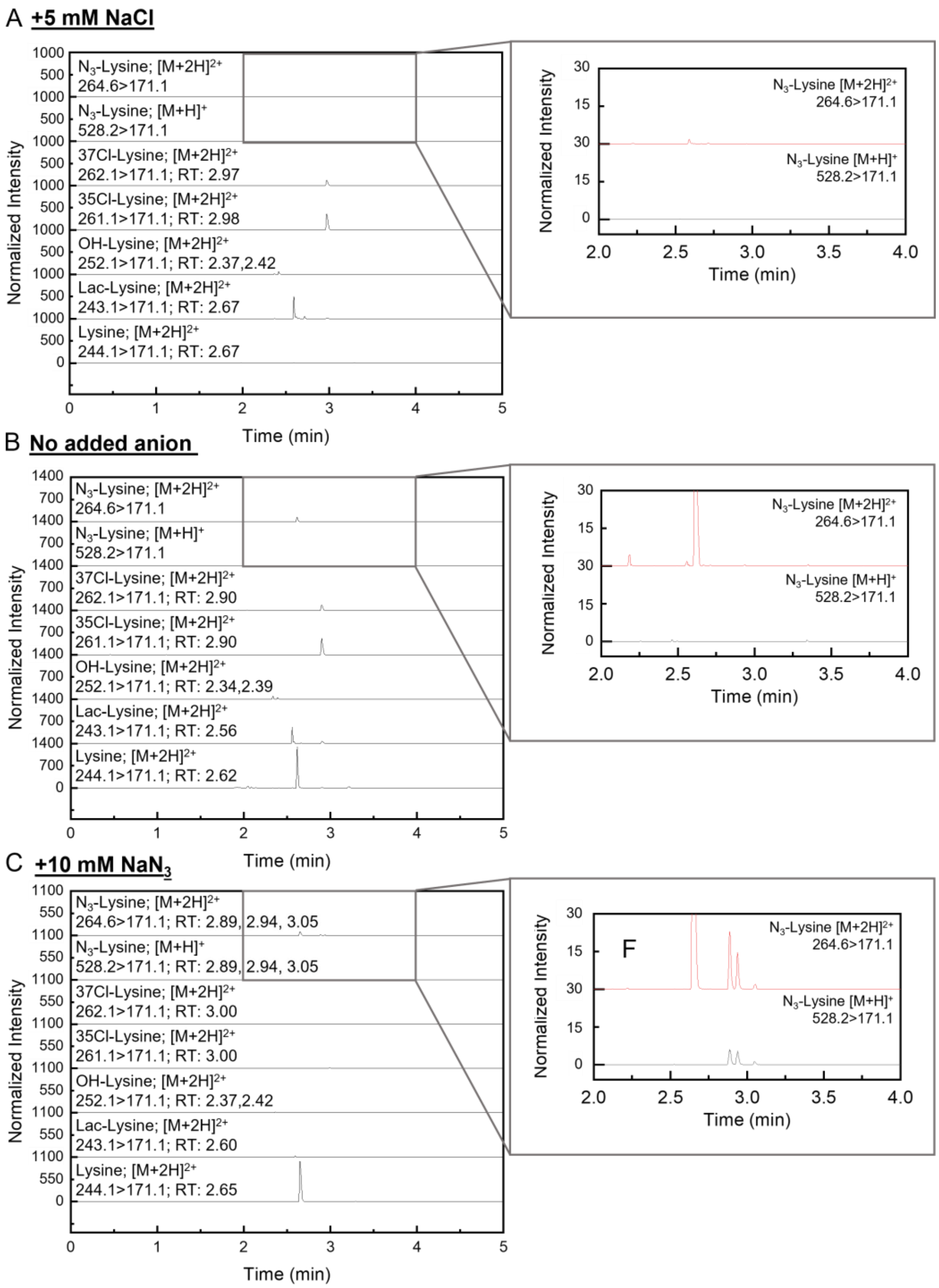


**Figure S7.** UPLC-MRM-MS/MS chromatograms of azidated lysine and other analytes. Reaction samples were prepared with 50 μM BesD, 5 mM 2OG, 1 mM sodium ascorbate, 100 μM $(NH_4)_2Fe(SO_4)_2$, 200 μM lysine in 100 mM HEPES (pH = 7.5) with either **(A)** 5 mM NaCl, **(B)** no added exogenous anion, or **(C)** 10 mM NaBr. Zoomed chromatograms of brominated lysine product channels **(D-F)** show peaks for brominated products only appear when bromide is directly added to the reaction. Accurate masses are confirmed in **Table S2**. All chromatograms shown have had intensity normalized to internal standard signal (L-Lysine(13C6, 99%; 15N2, 99%)·2HCl) intensity.

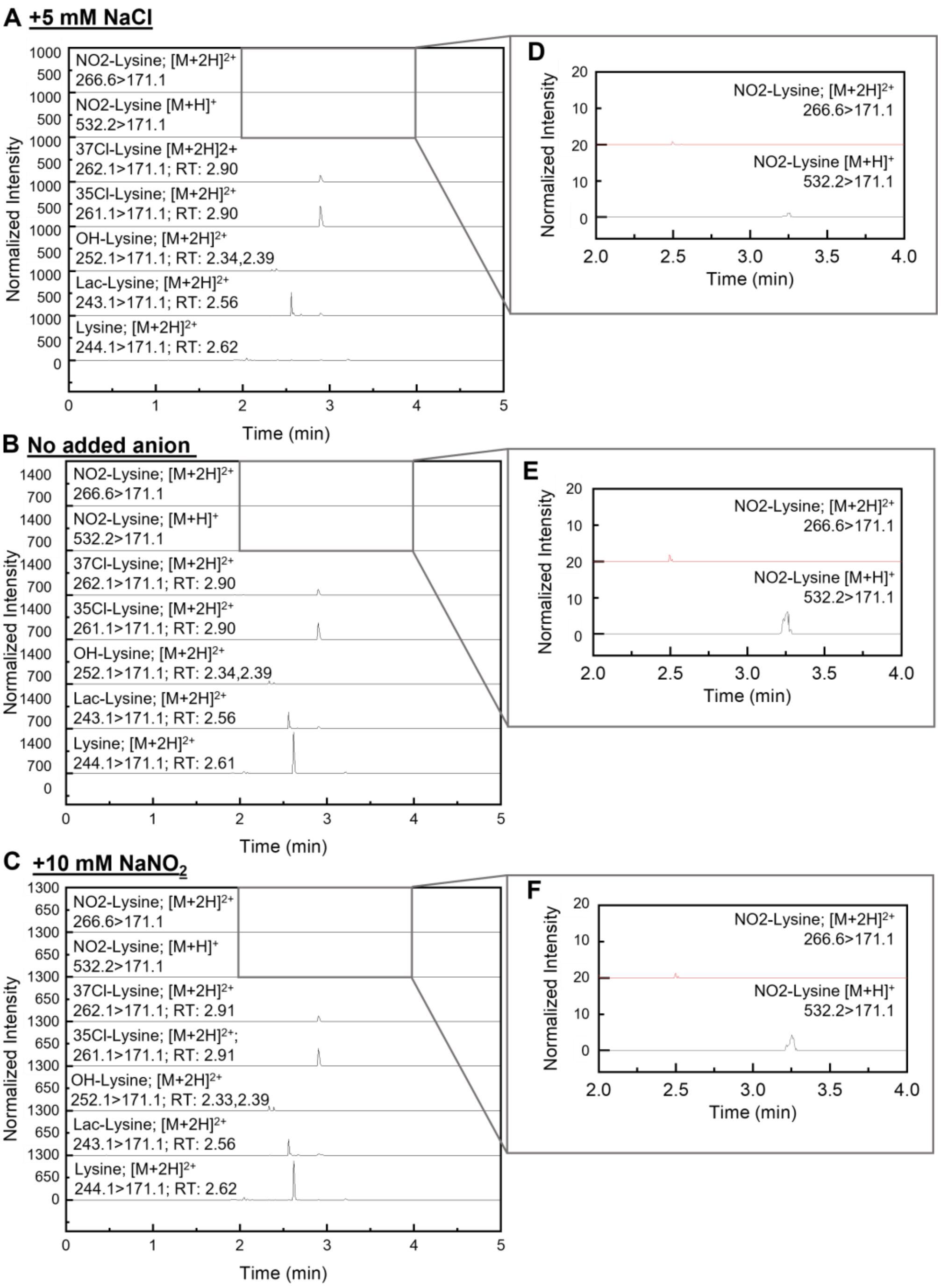


**Figure S8.** UPLC-MRM-MS/MS chromatograms of nitrated lysine and other analytes. Reaction samples were prepared with 50 μM BesD, 5 mM 2OG, 1 mM sodium ascorbate, 100 μM $(NH_4)_2Fe(SO_4)_2$, 200 μM lysine in 100 mM HEPES (pH = 7.5) with either **(A)** 5 mM NaCl, **(B)** no added exogenous anion, or **(C)** 10 mM $NaNO_2$. Zoomed chromatograms of nitrated lysine product channels **(D-F)** in area most likely to find derivatized lysine products indicate no peaks with masses corresponding to nitrated lysine. All chromatograms shown have had intensity normalized to internal standard signal (L-Lysine(13C6, 99%; 15N2, 99%)·2HCl) intensity.

### References


(1) Waterhouse, A.; Bertoni, M.; Bienert, S.; Studer, G.; Tauriello, G.; Gumienny, R.; Heer, F. T.; De Beer, T. A. P.; Rempfer, C.; Bordoli, L.; et al. SWISS-MODEL: Homology Modelling of Protein Structures and Complexes. *Nucleic Acids Res.* **2018**, *46* (W1), W296–W303. https://doi.org/10.1093/nar/gky427.

(2) Neugebauer, M. E.; Sumida, K. H.; Pelton, J. G.; McMurry, J. L.; Marchand, J. A.; Chang, M. C. Y. A Family of Radical Halogenases for the Engineering of Amino-Acid-Based Products. *Nat. Chem. Biol.* **2019**, *15* (10), 1009–1016. https://doi.org/10.1038/s41589-019-0355-x.

(3) Bienert, S.; Waterhouse, A.; Beer, T. A. P. De; Tauriello, G.; Studer, G.; Bordoli, L.; Schwede, T. The SWISS-MODEL Repository — New Features and Functionality. *Nucleic Acids Res.* **2017**, *45*, 313–319. https://doi.org/10.1093/nar/gkw1132.

(4) Bertoni, M.; Kiefer, F.; Biasini, M.; Bordoli, L.; Schwede, T. Modeling Protein Quaternary Structure of Homo- and Hetero- Oligomers beyond Binary Interactions by Homology. *Sci. Rep.* **2017**, No. March, 1–15. https://doi.org/10.1038/s41598-017-09654-8.

(5) Guex, N.; Peitsch, M. C.; Schwede, T. Automated Comparative Protein Structure Modeling with SWISS-MODEL and Swiss-PdbViewer: A Historical Perspective. *Electrophoresis* **2009**, *30*, S162–S173.

(6) Studer, G.; Rempfer, C.; Waterhouse, A. M.; Gumienny, R.; Haas, J.; Schwede, T. QMEANDisCo — Distance Constraints Applied on Model Quality Estimation. *Bioinformatics* **2020**, *36* (6), 1765–1771. https://doi.org/10.1093/bioinformatics/btz828.

(7) Gordon, J. C.; Myers, J. B.; Folta, T.; Shoja, V.; Heath, L. S.; Onufriev, A. H ++ : A Server for Estimating p K a s and Adding Missing Hydrogens to Macromolecules. *Nucleic Acids Res.* **2005**, *33*, 368–371. https://doi.org/10.1093/nar/gki464.

(8) Anandakrishnan, R.; Aguilar, B.; Onufriev, A. V. H++ 3.0 : Automating PK Prediction and the Preparation of Biomolecular Structures for Atomistic Molecular Modeling and Simulations. *Nucleic Acids Res.* **2012**, *40*, 537–541. https://doi.org/10.1093/nar/gks375.

(9) Myers, J.; Grothaus, G.; Narayanan, S.; Onufriev, A. A Simple Clustering Algorithm Can Be Accurate Enough for Use in Calculations of PKs in Macromolecules. *PROTEINS Struct. Funct. Bioinforma.* **2006**, *63*, 928–938. https://doi.org/10.1002/prot.

(10) Case, D. A.; Belfon, K.; Ben-Shalom, I. Y.; Brozell, S. R.; Cerutti, D. S.; Cheatham, T. E. I.; Cruzeiro, V. W. D.; Darden, T. A.; Duke, R. E.; Giambasu, G.; et al. AMBER 2020. University of California: San Francisco 2020.

(11) Tian, C.; Kasavajhala, K.; Belfon, K. A. A.; Raguette, L.; Huang, H.; Migues, A.; Bickel, J.; Wang, Y.; Pincay, J.; Wu, Q.; et al. Ff19SB: Amino-Acid-Specific Protein Backbone Parameters Trained against Quantum Mechanics Energy Surfaces in Solution. *J. Chem. Theory Comput.* **2020**, *16*, 528–552. https://doi.org/10.1021/acs.jctc.9b00591.

(12) Wang, J.; Wolf, R. M.; Caldwell, J. W.; Kollman, P. A.; Case, D. A. Development and Testing of a General AMBER Force Field. *J. Comput. Chem.* **2004**, *25*, 1157–1174.

(13) Li, Z.; Song, L. F.; Li, P.; Merz, K. M. Systematic Parametrization of Divalent Metal Ions for

the OPC3, OPC, TIP3P-FB, and TIP4P-FB Water Models. *J. Chem. Theory Comput.* **2020**, *16*, 4429−4442. https://doi.org/10.1021/acs.jctc.0c00194.

(14) Wang, J.; Wang, W.; Kollman, P. A.; Case, D. A. Automatic Atom Type and Bond Type Perception in Molecular Mechanical Calculations. *J. Mol. Graph. Model.* **2006**, *25*, 247–260.

(15) Izadi, S.; Anandakrishnan, R.; Onufriev, A. V. Building Water Models : A Different Approach. *J. Phys. Chem. Lett.* **2014**, *5*, 3863−3871.

(16) Horn, A. H. C. A Consistent Force Field Parameter Set for Zwitterionic Amino Acid Residues. *J. Mol. Model.* **2014**, *20*. https://doi.org/10.1007/s00894-014-2478-z.

(17) Li, P.; Merz, K. M. MCPB.Py: A Python Based Metal Center Parameter Builder. *J. Chem. Inf. Model.* **2016**, *56* (4), 599–604. https://doi.org/10.1021/acs.jcim.5b00674.

(18) Case, D. A.; Aktulga, H. M.; Belfon, K.; Ben-Shalom, I. Y.; Brozell, S. R.; Cerutti, D. S.; Cheatham, T. E. I.; Cisneros, G. A.; Cruzeiro, V. W. D.; Darden, T. A.; et al. Amber 2021. University of California, San Francisco 2021.

(19) Becke, A. D. Density-Functional Thermochemistry. III. The Role of Exact Exchange. **2013**, *5648* (August 1998).

(20) Wilk, L.; Nusair, M. Accurate Spin-Dependent Electron Liquid Correlation Energies for Local Spin Density Calculations: A Critical Analysis1. **1980**.

(21) Lee, C.; Yang, W.; Parr, R. G. Development of the Colic-Salvetti Correlation-Energy into a Functional of the Electron Density. *Phys. Rev. B* **1988**, *37* (2), 785–789.

(22) Roe, D. R.; Cheatham, T. E. PTRAJ and CPPTRAJ: Software for Processing and Analysis of Molecular Dynamics Trajectory Data. *J. Chem. Theory Comput.* **2013**, *9*, 3084−3095.

(23) Nicholls, A. Confidence Limits , Error Bars and Method Comparison in Molecular Modeling . Part 1 : The Calculation of Confidence Intervals. **2014**, 887–918. https://doi.org/10.1007/s10822-014-9753-z.

(24) Neese, F. The ORCA Program System. **2012**, *2* (February), 73–78. https://doi.org/10.1002/wcms.81.

(25) Neese, F. Software Update: The ORCA Program System — Version 5 . 0. *Wiley Interdiscip. Rev. Comput. Mol. Sci.* **2022**, 1–15. https://doi.org/10.1002/wcms.1606.

(26) Grimme, S.; Ehrlich, S.; Goerigk, L. Effect of the Damping Function in Dispersion Corrected Density Functional Theory. **2011**, No. Sfb 858. https://doi.org/10.1002/jcc.

(27) Schäfer, A.; Horn, H.; Ahlrichs, R. Fully Optimized Contracted Gaussian Basis Sets for Atoms Li to Kr. *J. Chem. Phys.* **1992**, *97* (4), 2571–2577. https://doi.org/10.1063/1.463096.

(28) Schafer, A.; Huber, C.; Ahlrichs, R. Fully Optimized Contracted Gaussian Basis Sets of Triple Zeta Valence Quality for Atoms Li to Kr. **2020**, *5829* (November 1993).

(29) Phys, J. C.; Grimme, S.; Antony, J.; Ehrlich, S. Parametrization of Density Functional Dispersion Correction ( DFT-D ) for the 94 Elements H-Pu Dispersion Correction „ DFT-D … for the 94 Elements H-Pu. **2019**, *154104* (2010). https://doi.org/10.1063/1.3382344.

(30) Smith, D. G. A.; Burns, L. A.; Patkowski, K.; Sherrill, C. D. Revised Damping Parameters for the D3 Dispersion Correction to Density Functional Theory. **2016**. https://doi.org/10.1021/acs.jpclett.6b00780.

(31) Weigend, F. Accurate Coulomb-Fitting Basis Sets for H to Rn. *Phys. Chem. Chem. Phys.* **2006**, *8* (9), 1057–1065. https://doi.org/10.1039/b515623h.

(32) Lu, T.; Chen, F. Multiwfn: A Multifunctional Wavefunction Analyzer. *J. Comput. Chem.* **2012**, *33* (5), 580–592. https://doi.org/10.1002/jcc.22885.

(33) Zhang, J.; Lu, T. Efficient Evaluation of Electrostatic Potential with Computerized Optimized Code. *Phys. Chem. Chem. Phys.* **2021**, *23* (36), 20323–20328. https://doi.org/10.1039/d1cp02805g.

(34) Lu, T.; Chen, F. Quantitative Analysis of Molecular Surface Based on Improved Marching Tetrahedra Algorithm. *J. Mol. Graph. Model.* **2012**, *38*, 314–323. https://doi.org/10.1016/j.jmgm.2012.07.004.

(35) Humphrey, W.; Dalke, A.; Schulten, K. VMD - Visual Molecular Dynamics. *J. Molec. Graph.* **1996**, *14*, 33–38.

(36) Wilson, R. H.; Chatterjee, S.; Smithwick, E. R.; Dalluge, J. J.; Bhagi-, A. Role of a Secondary Coordination Sphere Residue in Halogenation Catalysis of Non-Heme Iron Enzymes. *ACS Catal.* **2022**, *12*, 10913−10924. https://doi.org/10.1021/acscatal.2c00954.
